# Human Herpes Virus-8 Oral Shedding Heterogeneity is Due to Varying Rates of Reactivation from Latency and Immune Containment

**DOI:** 10.1101/2024.11.26.625350

**Authors:** David A Swan, Elizabeth M Krantz, Catherine Byrne, Fred Okuku, Janet Nankoma, Innocent Mutyaba, Warren Phipps, Joshua T. Schiffer

**Affiliations:** Vaccine and Infectious Diseases Division, Fred Hutchinson Cancer Center, Seattle, USA; Uganda Cancer Institute, Kampala, Uganda; Department of Medicine, University of Washington, Seattle, USA

## Abstract

Human herpesvirus-8 (HHV-8) is a gamma herpesvirus linked to the development of Kaposi sarcoma (KS). KS is more common in persons living with HIV (PLWH), but endemic KS in HIV-negative individuals is also common in sub-Saharan Africa. HHV-8 shedding occurs in the oral mucosa and is likely responsible for transmission. The mechanistic drivers of different HHV-8 shedding patterns in infected individuals are unknown. We applied stochastic mathematical models to a longitudinal study of HHV-8 oral shedding in 295 individuals in Uganda who were monitored daily with oral swabs. Participants were divided into four groups based on whether they were HIV-negative or positive as well as KS-negative or positive. In all groups, we observed a wide variance of shedding patterns, including no shedding, episodic low viral load shedding, and persistent high viral load shedding. Our model closely replicates patterns in individual data and attributes higher shedding rates to increased rates of viral reactivation, and lower median viral load values to more rapid and effective engagement of cytolytic immune responses. Our model provides a framework for understanding different shedding patterns observed in individuals with HHV-8 infection.

**Keypoints:** HHV8 shedding rate is mosty determined by rate of reactivation from latency while viral loads is mostly dteremined by peripheral immune responses.

DAS performed all mathematical modeling and editied the paper; EMK performed statistical analysis and edited the paper; CB assisted with modeling; FO, JN and IM designed and implemented the clinical protocols; WP designed and implemented the clinical protocols and edited the paper; JTS conceived the study and write the paper.

## Introduction

Human herpesvirus-8 (HHV-8) is a gamma herpesvirus that was discovered in 1994 as the causative agent of Kaposi Sarcoma (KS),^1^ a hyperproliferative endothelium cancer most often observed in individuals with HIV.^2^ Endemic KS occurs in HHV-8-positive, HIV-negative individuals and is common in sub-Saharan Africa.^3, 4^ Yet, most individuals with HHV-8, even those who shed the virus orally frequently and at high viral loads, do not develop KS.^5–8^ The link between oral shedding and transmission, as well as KS tumorigenesis, remains unclear.^5, 9–11^

HHV-8 and Epstein-Barr virus (EBV), the other human gamma herpesvirus, are similar in many aspects. Both establish latency within B cells, while HHV-8 may become latent in epithelial and endothelial cells allowing the virus to persist for the lifetime of the human host in a non-replicative state.^12, 13^ HHV-8 and EBV have been implicated in oncogenic disorders of B cells, including Multicentric Castleman’s disease (MCD)^14–17^ and primary effusion lymphoma (PEL)^18, 19^ for HHV-8, and several B cell lymphomas for EBV.^20^ HHV-8 and EBV establish lytic infection in both endothelial and epithelial cells.^21^ Viral replication in oral epithelial cells leads to substantial amplification of viral DNA and may facilitate viral transmission.^5, 22^

Viral shedding kinetics provide a key window into host-virus interactions and differ significantly between different human viral pathogens, as well as the immune status of infected individuals. Mathematical models are a critical tool for the analysis of longitudinal viral load data and account for the non-linearity inherent to this data.^23^ Models have been applied to capture the timing and intensity of immune responses in tissue,^24^ to compare different severity of disease among multiple infected individuals,^25^ and to optimize therapeutic and vaccination approaches.^26^ For human herpes viruses including herpes simplex virus-2 (HSV-2),^27, 28^ cytomegalovirus (CMV),^29^ and EBV,^13, 30^ stochastic models are required given the episodic and unpredictable nature of shedding within chronically infected individuals. While these models cannot be used to precisely recapitulate an individual’s shedding trajectories, they can capture patterns of shedding using summary statistics such as median and variability of viral load, episode rate, expansion and clearance slope, and peak viral loads.^31^ While shedding kinetics differ markedly among human herpes viruses, a commonality is that critical immune responses occur in micro-environments and often over narrow time intervals.^5, 30, 32, 33^ Here, we develop the first mathematical models to describe HHV-8 shedding data in the oral cavity to understand observed shedding patterns in a Ugandan cohort of individuals with and without HIV infection and with and without KS.^11^

## Methods

### Mathematical Model of HHV-8 Oral Shedding

We developed a stochastic mathematical model to understand the basis of observed heterogeneous HHV-8 shedding patterns in the oral mucosa. Our model focuses on key parameters observed in our recent EBV model including rate of viral reactivation from latency and effectiveness of the peripheral immune responses in eliminating infected cells.^30^ Our model also assumes multiple micro-regions to simulate the oral cavity based on analysis done in previous modeling of EBV.^30^ Each region is assumed to have a certain probability of viral reactivation and a separate immunologic response **(Fig 1)**.

**Figure 1:**
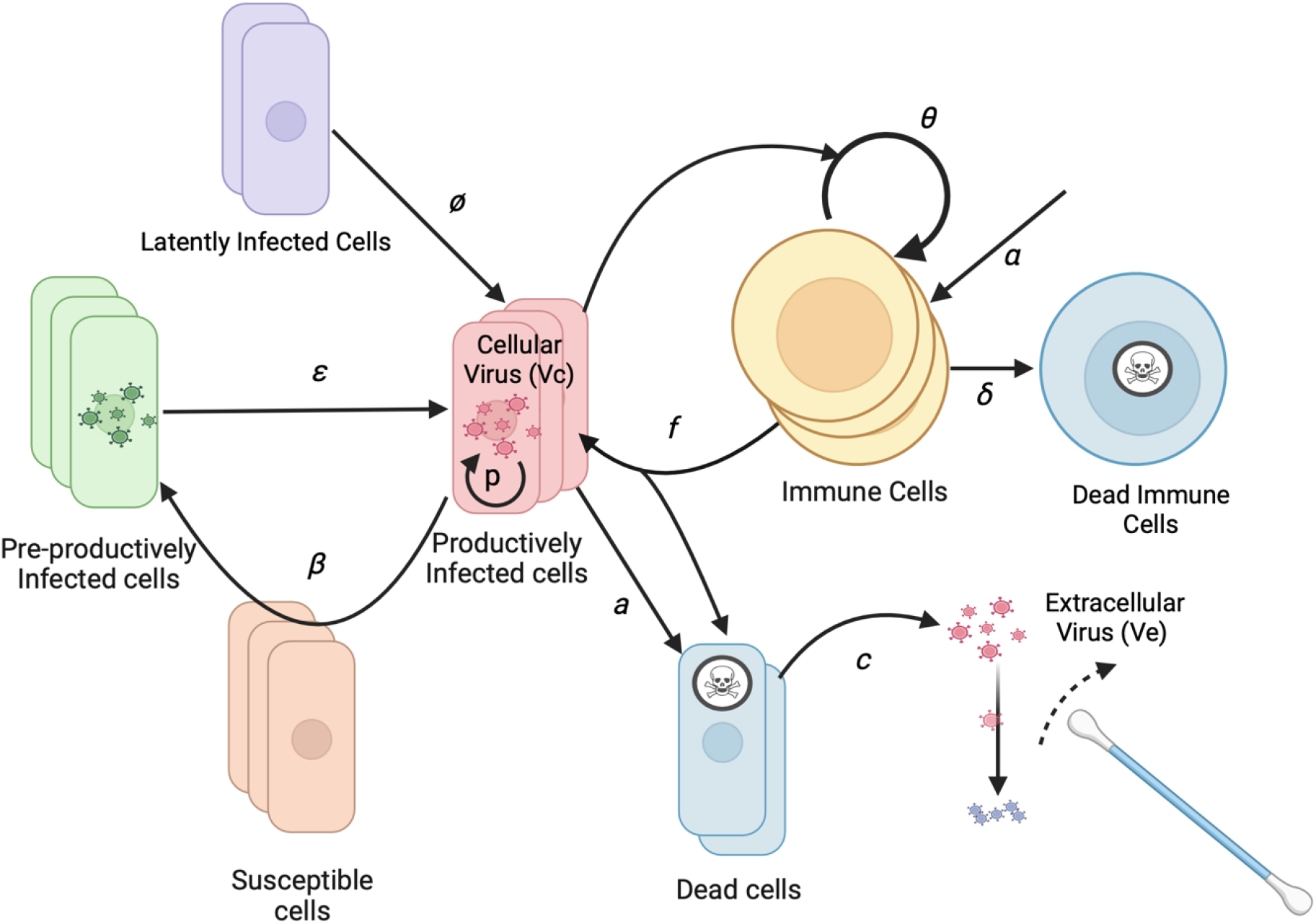
HHV-8 Mathematical Model Schematic for Infection within the Oral Mucosa. Basic model assumptions are that latently infected cells reactivate at rate ϕ. Infection spreads from cell to cell with infectivity β. Cells exit an initial non-productive eclipse phase at rate ε. Infected cells die after becoming lytic at rate *a*. Infected cells produce virus at rate *p*. Cytolytic immune cells expand at rate θ that becomes half maximal when *r50* cells are infected and kill infected cells at rate *f*, and are produced independently of infected cells at rate α and die at rate δ. Cell-associated virus converts to cell-free virus upon cell death and then decays at rate *c*. Cell-free virus is detected with oral swabs. Multiple regions can produce virus concurrently.

To limit model parameters, we assigned an infectivity to each infected cell (I), assuming a certain rate of viral production, rather than modeling the infectivity of each virion. The model includes a reactivation rate ϕ which dictates how frequently latently infected cells become lytic. When lytic infection is initiated, an infected cell begins producing virus and infecting adjacent cells. While the reactivating cell may be a B cell, B cells and reseeding of the latently infected cell reservoir are not explicitly included as model variables due to the lack of available data to parameterize the dynamics of these cell types separately. After an eclipse period (E) of average duration 1/ε, the infected cell begins producing cell-associated virus (Vc) at a rate of *p* virions per day. The infection spreads to additional cells in the region based on infectivity parameter β.

As the population of infected cells in a region increases, the population of local HHV-8 specific T cells (T) may increase and kill infected cells at a rate *f*. Infected cells also die (or cease viral production) at a rate *a*. At that same rate, cell-associated virus is released from the cell and becomes cell-free virus (Ve) which is captured during daily swabbing of the oral cavity. Cell-free virus then decays at a rate *c*. The T cell population in each region grows at a maximal rate of θ. The rate is modulated by the number of infected cells in the regions with it being half maximal at a level of *r50* cells. T cells also arrive at a rate α into each region and die at a rate of δ. The initial T cell population in each region is randomly seeded and then allowed to equilibrate by running the model for a year before fitting to data.

The stochastic differential equations for the model are shown below. Each region is represented using the subscript j. The major compartments are E_j_ for infected cells in the eclipse phase, I_j_ for productively infected cells, T_j_ for T cells, Vc_j_ for cell associated virus and Ve_j_ for extra-cellular virus. Vetot is the amount of virus available for oral swabbing at each time step and is the sum of all regions.

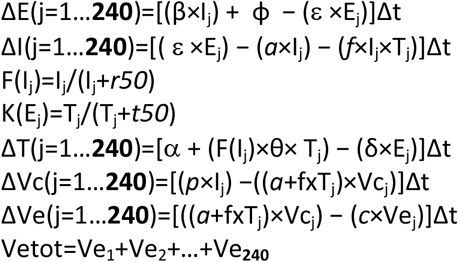

The model does not include viral spread between micro-environments. When multiple regions are involved during an episode, it is therefore due to multiple stochastic sites of reactivation from latency.

We also tested a model where virions initiate the spread of infection among cells in each region and where cells in one region can infect cells in adjacent regions. The latter did not fit the data as well despite its added complexity (results not shown).

### Cohort data

We fit the model to data from an observational, prospective cohort study of Ugandan adults (age ≥18 years) who were enrolled between October 2007 and May 2010. Individuals were from four groups: HIV-1 seropositive individuals without KS (HIV+/KS-); HIV-1 seronegative individuals without KS (HIV-/KS-); HIV-1 seropositive individuals with KS (HIV+/KS+); and HIV-1 seronegative individuals with KS (HIV-/KS+). HIV+/KS-participants who reported ART use at the time of enrollment were excluded. HIV+/KS+ participants were eligible despite ART use. At the time of original study, ART was initiated based on CD4+ T cell levels and universal test and treat policies were not yet established. Participants from all arms were followed for at least 4 weeks, with one “session” of data collection consisting of 28 days of daily home oral swab collection. This monthly cycle of sample collection was repeated every three months for Arm A (HIV+/KS-) participants (up to two years) and for Arm C (HIV+/KS+) participants (up to one year).^34^

Oral swab samples were evaluated for KSHV DNA by a quantitative, high-throughput fluorescent-probe-based real-time polymerase chain reaction (PCR) assay (TaqMan assay, Applied Biosystems, Foster City, CA) of the ORF73 gene at the UCI-Fred Hutch Cancer Centre Laboratory in Kampala, Uganda as described previously.^9, 10^ Samples with >150 copies per mL of KSHV DNA were considered positive.^35^

This study was approved by the Makerere University Research and Ethics Committee, the Fred Hutch Cancer Center Institutional Review Board, and the University of Washington IRB. The data has been described in our prior work.^34^

### Model Fitting

Our model simulations are intended to match data from 295 individuals enrolled in the natural history cohort study who had at least one positive oral sample during the initial 4-week sampling period. Rather than fit to individual viral trajectories, we attempted to reproduce three summary measures from each person: shedding rate (percentage of swabs positive for HHV8 DNA by PCR), median log HHV8 viral load per positive swab and peak log HHV8 viral load. To match the study protocol, the simulation data was sampled daily for 28 consecutive simulated days.

Parameters for the models were fixed when possible using values from the literature, or fitted to match the observed data. All fixed and fitted parameters are shown in **Table 1** together with their values and references.

**Table 1:** Model Parameters.

| Parameter | Symbol | Value | Reference |
| --- | --- | --- | --- |
| Max T cell growth rate | $\theta$ | 3 / day | 28, 36 |
| Eclipse period | $\varepsilon$ | 0.85 days | 37 |
| T cell birth rate | $\alpha$ | 0.312 / day | *Fitting (range 0.1-0.5) |
| T cell death rate | $\delta$ | 0.001 / day | 28 |
| Infected cells for half-maximal T cell growth | $r_{50}$ | 300 cells | *Fitting (range 2-3.5 log10) |
| Viral clearance rate | $c$ | 6 / day | *Fitting (range 5-20) |
| Infected cell death rate | $a$ | mean=1.48 / day<br>sd=0.01 | Fitting (range 0.5-2.5) |
| Cell infectivity | $\beta$ | mean=8.2 cells / day<br>sd=1.4 | Fitting (range 1-20) |
| Log Viral production | Log $p$ | mean=5.86 log10<br>viruses / cell / day<br>sd=1.1 log10 | Fitting (range 3.5-7 log10) |
| T cell killing rate | $f$ | mean=23.5 cells / day<br>sd=24 | Fitting (range -2 – 2 log10) |
| Reactivation rate | $\phi$ | mean=1.08 cells / day<br>sd=1.7 | Fitting (range -1 – 2 log10) |

\*Some of the fixed parameters were initially fitted and later fixed as their final best ranges proved to be quite narrow. These included the birth rate of T cells, the level of infected cells at which T cell growth is half maximal, the T cell exhaustion rate, and viral clearance rates. Limiting the set of fitted parameters allowed more in-depth exploration of their parameter space.

Initially, fitted parameters were drawn from wide, primarily log-linear distributions. One thousand sets of simulations were run, each with a different set of values for the fitted parameters. Adaptive immunity levels were established by running the simulation for one year and measuring T cell levels in each region. Since the simulations were run stochastically, 10 runs were performed for each unique set of parameters. The starting T cell distributions were used in each run to save computational time, but sampling was delayed until day 150 to allow for on-going episodes to be captured. The median and peak log HHV-8 viral loads and shedding rates were averaged across runs. Each of the 1,000 simulations was scored against the data for that individual. The score was composed of the average of the percent error (the absolute difference between simulated and target values as a percentage of the target value) for each of the three measures. The parameter values for the top 100 runs were used to determine a posterior distribution for each parameter which were defined using their values for mean and standard deviation. Those distributions were used to draw 1,000 new parameter combinations and the process was repeated. After two such rounds of simulations to narrow distributions, the mean values of each fitted parameter were recorded. These were then used in a set of follow-on simulations (10 per participant) with the average simulated metrics used to determine concordance between simulated and actual data across all participants.

## Results

### Cohort data

Thirty-six (29%) HIV+/KS-participants, 16 (21%) HIV-/KS-, 56 (74%) HIV+/KS+ and 9 (50%) HIV-/KS+ participants had HHV-8 detected in at least one oral swab. Data from these individuals was used for model fitting. As we described in an earlier analysis of oral HHV-8 shedding in the cohort study, shedding rates and median viral load varied substantially across individuals in all four groups, and were correlated with one another.^34^

### Model simulations recapitulate observed variability in HHV-8 shedding

Mathematical model output showed high correlation with the observed shedding patterns across all individuals with positive swabs during the study period as shown in **Fig 2**. The concordance correlation coefficient (CCC), a measure of correlation and agreement between simulated and observed data was 0.988 for shedding rate **(Fig 2a)**, 0.915 for median log HHV8 **(Fig 2b)** and 0.922 for peak viral load **(Fig 2c)**. Modeled peak viral load slightly exceeded observed low peaked episodes and were often slightly lower than the high observed high peaks **(Fig 2C)**. Pearson correlation coeficients were also high for each metric **(Fig 2a-c)**. Each parameter combination was also scored using the average percentage error between each individual’s shedding metrics and simulated output as described in the fitting section. Averaging the scores for the top 10 parameter combinations per participant across the three categories, the average percent error was 3.7%, again indicating that the model output is very close to the cohort data.

**Figure 2:**
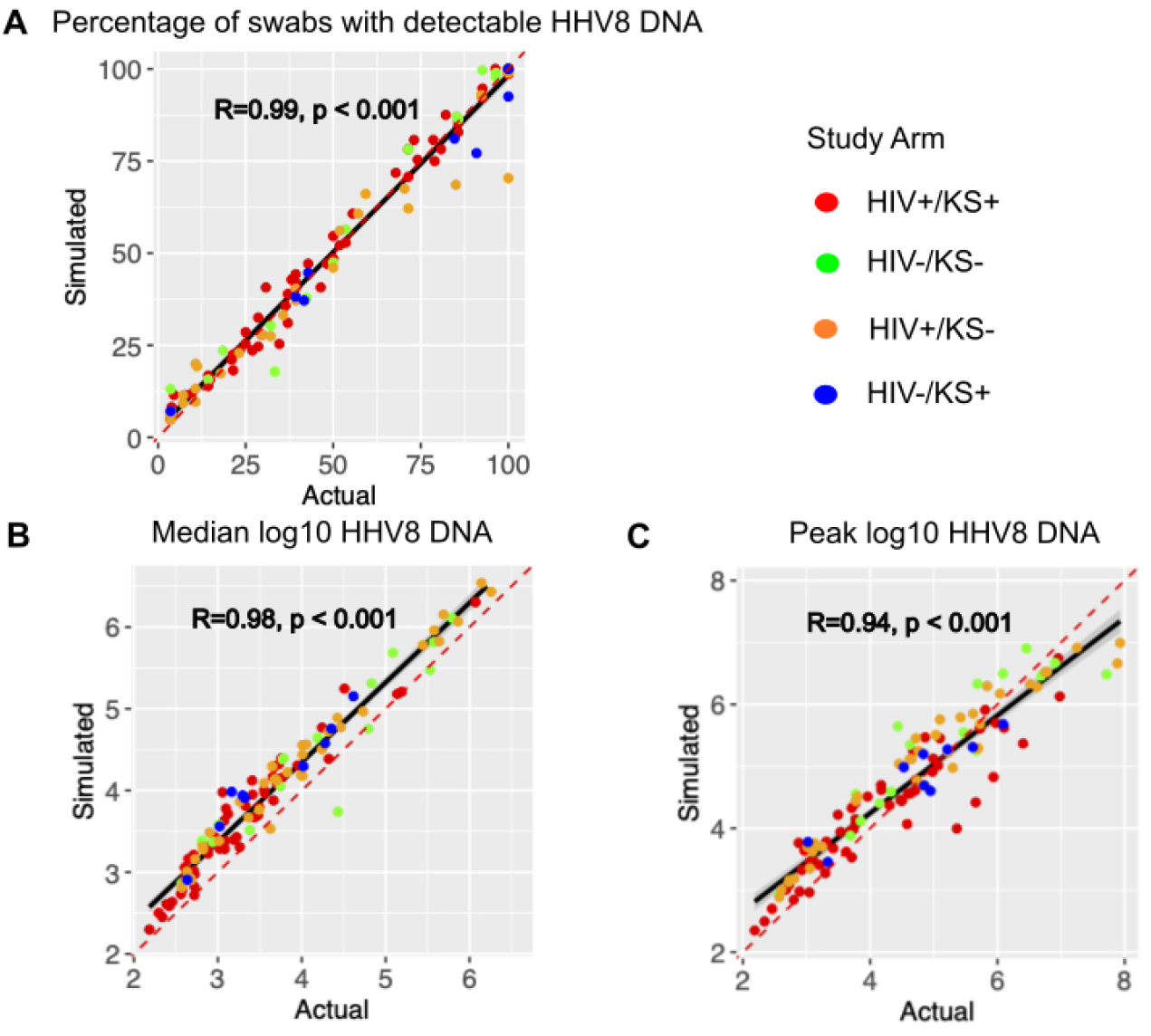
Mathematical model output reproduces individual shedding metrics. Actual (x-axis) versus simulated (y-axis) for (a) percentage of positive swabs, (b) median log10 HHV-8 DNA copies, and (c) peak log10 HHV-8 copies. Each dot is a study participant. The red dotted line is the line of perfect concordance. The solid black line is a concordance line of actual versus simulated data. Participants are colored by study arm as in the legend. R is the Pearson correlation coefficient.

We previously demonstrated in the cohort study that HHV-8 shedding occurs along a gradient.^34^ On one extreme, certain individuals shed at a rate exceeding 80%, typically at high median viral loads >10^5^ copies/swab which may vary over time over a range of several logs. Other infected people shed episodically at a low rate (<20%), typically at much lower median viral loads (<10^3^ copies/swab), and rarely with a single sample exceeding 10^5^ copies/swab. Many individuals have shedding patterns falling between those two extremes.

HHV-8 viral load traces were plotted from 3 different representative participants in each of three different shedding categories: infrequent shedding (1 or 2 positive swabs during the 28 days, **Fig 3a**), medium shedding (∼30-70% of the swabs being positive, **Fig 3c**) and constant shedding (all swabs positive, **Fig 3e**). For each participant, three separate simulations were run using the best mean values of their individually estimated parameters. The simulation run plots show patterns compatible with observed shedding **(Fig 3b,d,Ff** despite the stochastic nature of the model output.

**Figure 3.**
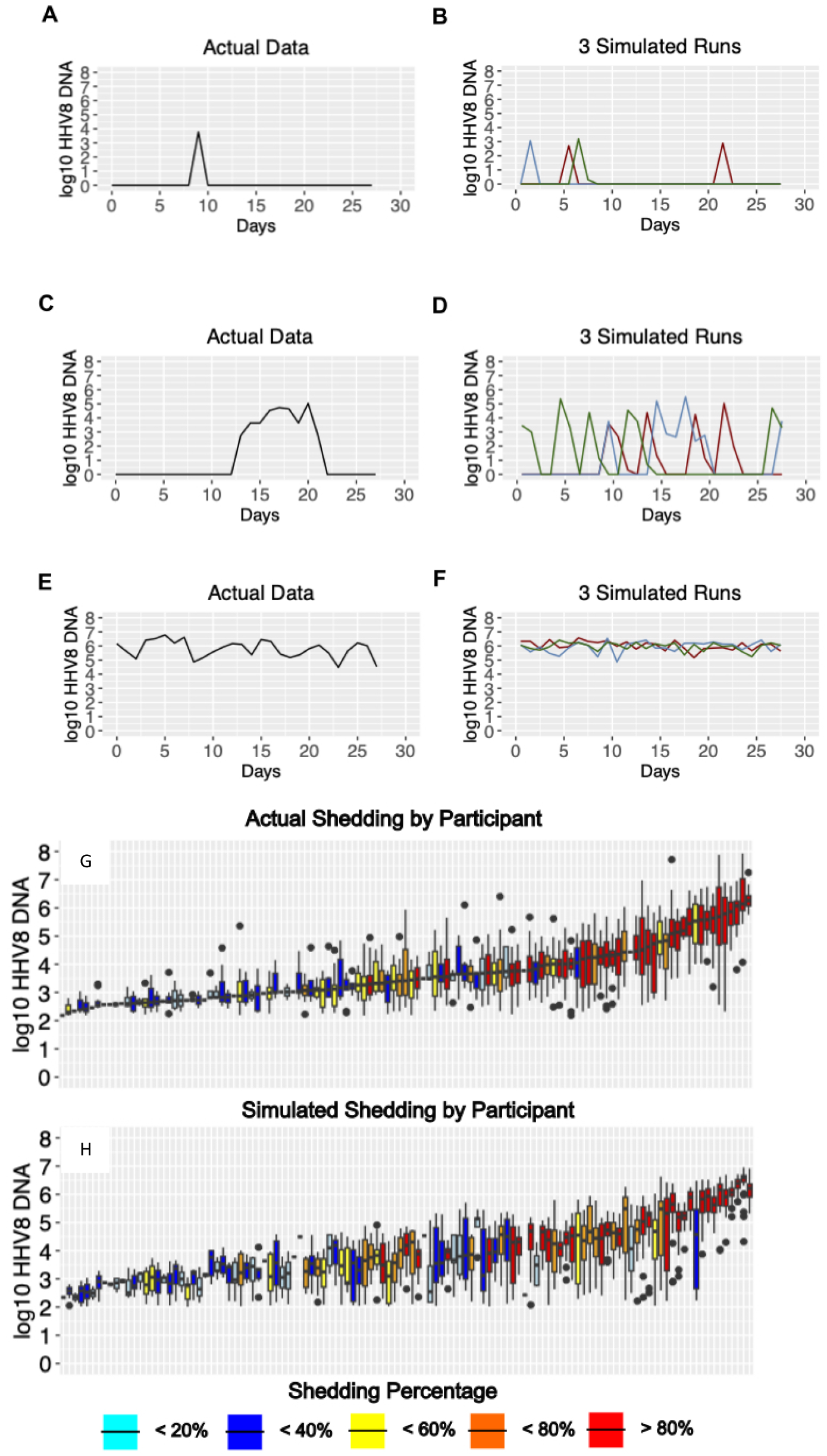
Model output matches three observed HHV8 shedding types. Study data from three participants showing (a) rare episodic shedding, (c) more prolonged episodic shedding, and (e) constant higher viral load shedding is matched in a general fashion by model simulations in (b), (d), and (f) respectively. Low threshold curoff was 150 DNA copies per sample. (g,h) Distribution of oral HHV-8 viral loads for actual and simulated study participants. (g) Actual participant data sorted from left to right by median viral load and (h) simulated participant data matched to actual data in (g). Each boxplot shows data from a participant; boxes represent interquartile range, horizontal lines within boxes represent medians and whiskers extend to minimum and maximum values. Participants with only a median bar and no box had too few positive samples to generate an interquartile range. Participants are color coded according to shedding.

In the cohort study, we observed high variability in shedding rate in all 4 study groups. However, individuals with high shedding rates also tended to have higher median viral loads **(Fig 3g)**, a trend that was also observed in the simulated output **(Fig 3h)**.

### Mechanistic predictions of HHV-8 shedding rate and viral load

We next used our simulated output to determine how mathematical model parameter variability correlated with key shedding outcomes. We observed that a higher viral reactivation rate was more predictive of a participant’s HHV-8 shedding rate than other model parameter values **(Fig 4a, c, e)** whereas a higher viral per cell production rate and lower immune killing rate of infected cells was more predictive of higher median viral load **(Fig 4b, d, f)**. Overall, all three model inputs did have at least moderate impact on shedding rate and viral load.

**Figure 4:**
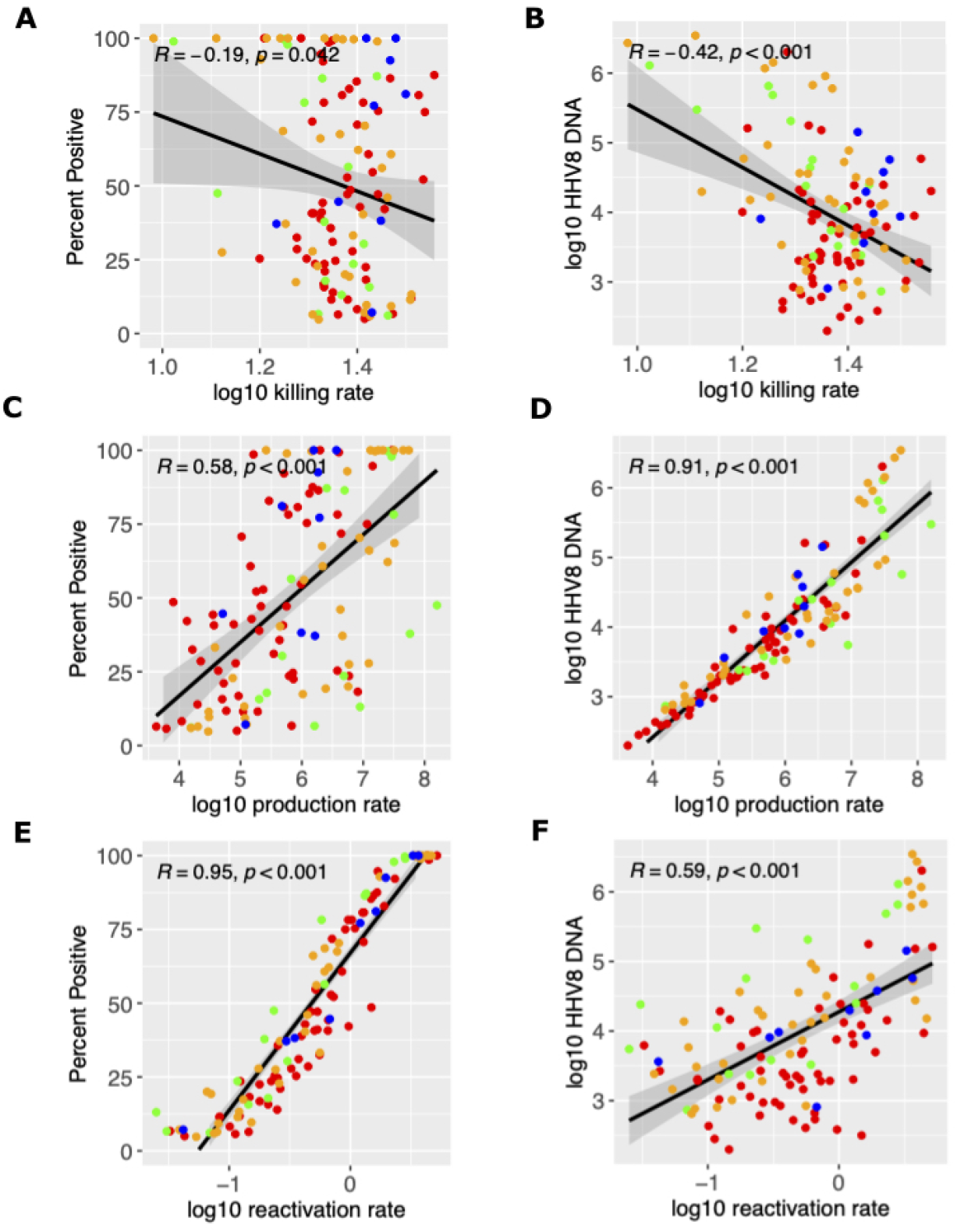
Mathematical model mechanistic predictors of shedding rate and median viral load. Possible determinants of outcomes are x-axes and include (a-b) log10 killing rate of infected cells, (c-d) log10 viral production rate per cell and and (e-f) log10 viral reactivation rate from latency. Y-axes are outcomes include (a,c,e) shedding rate and (b,d,f) median viral load. Each dot is a simulated participant.

### Parameter ranges according to HIV and KS status

We observed high variability among all parameter values for all study groups (**Fig 5)**. The average immune cell killing rate was slightly higher for the HIV−/KS+ groups than other groups **(Fig 5a)**. The mean per cell viral production rate was highest in HIV−/KS− individuals and lowest in HIV−/KS+ individuals **(Fig 5b)**. We also observed slightly higher HHV-8 reactivation rates in the HIV−/KS+ group versus the other groups **(Fig 5c)**.

**Figure 5:**
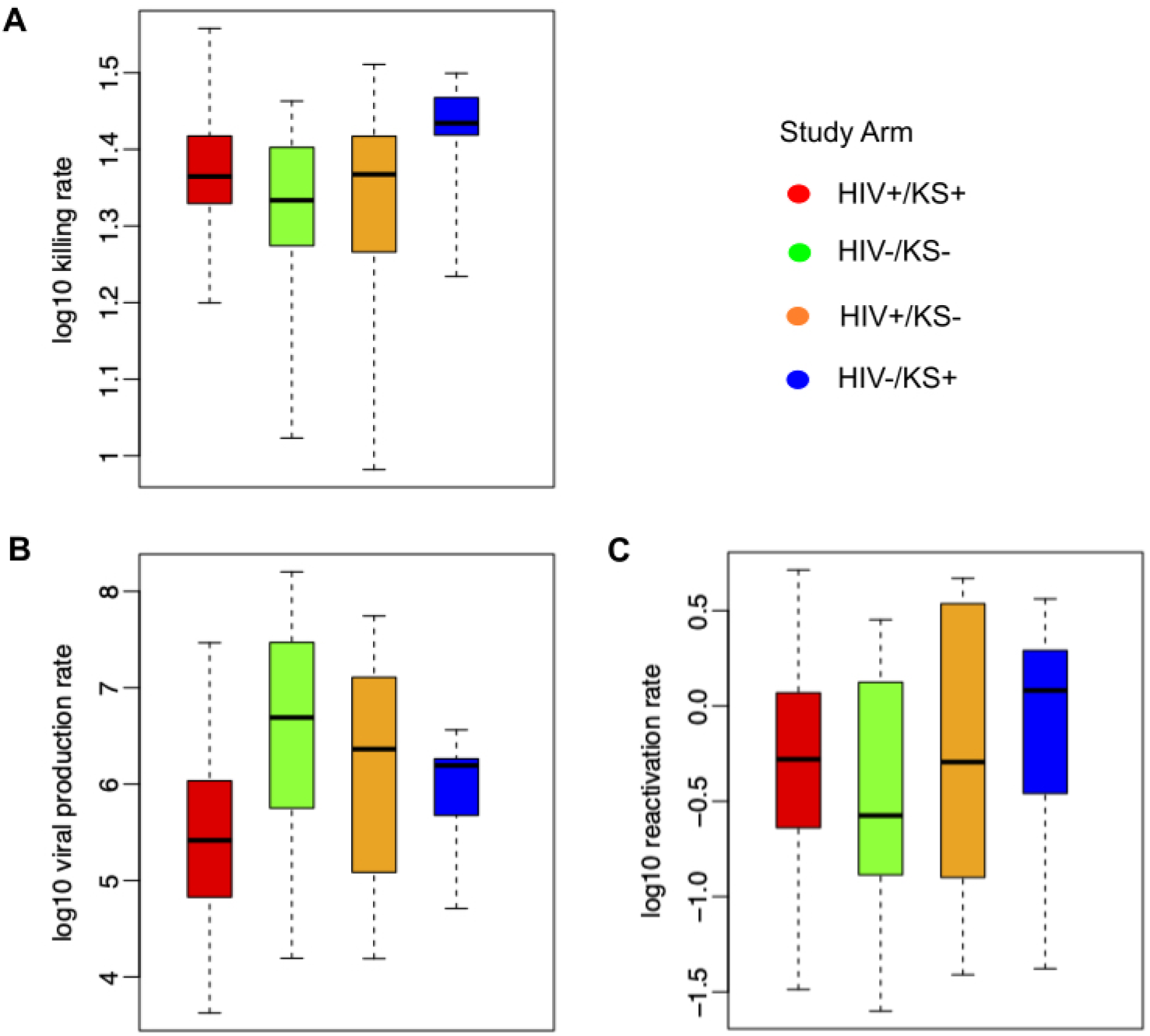
Mathematical model parameter ranges according to HIV and KS status. (a) Ranges of model-derived immune cell killing rates, (b) per cell viral production rates, and (c) latency viral reactivation rates. Each boxplot shows data from all participants in each group; boxes represent interquartile range, horizontal lines within boxes represent medians and whiskers extend to minimum and maximum values.

## Discussion

We used a mathematical model to capture HHV-8 shedding kinetics in a diverse cohort of individuals stratified by HIV-1 infection and KS status. The model’s stochastic output does not allow precise recapitulation of individual oral shedding patterns, but does reproduce key metrics of individuals in the cohort, including shedding rate, median viral load and peak viral load. The simulated data is therefore a close representation of the true data at the individual level and across the entire cohort.

Each human herpesvirus is notable for a specific pattern of shedding which differs according to numerous factors, including viral detection rate, episodicity versus persistence, viral expansion rate, viral clearance rate, median viral load, peak viral load and stability over various time intervals. We have applied mathematical models to most of these viruses to develop hypotheses regarding their observed shedding kinetic profile.^27–29, 32, 38^ We concistently identified that herpes virus shedding is ultimately driven by rates of viral reactivation from latency, rates of viral amplification in the lytic epithelial cell compartment, and pace of the antiviral immune response. A shared feature among viruses including HSV-1 and -2,^27, 28^ human herpes virus-6 (HHV-6),^32, 38^ CMV,^29, 39^ and EBV,^30^ is that variable observed shedding rates and duration of shedding events can be attributed to the latency reactivation rate and rate of peripheral immune control. Our current modeling suggests that HHV-8 appears to follow these trends as well.

Many specific features of HHV-8 shedding closely resemble EBV shedding, while differing significantly from the other human herpesviruses. Specifically, we observe an extremely wide range of HHV-8 shedding phenotypes, including rare and brief episodic shedding, more frequent and prolonged episodic shedding, and continual high viral load shedding.^30, 34^ Our model explains this heterogeneity according to variance in key biologic parameters. As with EBV, the rate of viral reactivation from latency is predicted to be highly associated with the observed shedding rate, while median viral load appears to have multi-factorial determinants including viral reactivation rate, rate of viral production from infected cells, and rate of killing by the immune system. In theory, each of these processes would be a viable therapeutic target, but even partial reduction in the rate of viral reactivation from latency might have an outsized effect on shedding and possible transmission likelihood.

There did not appear to be a clear set of shedding or model patterns that clearly differentiate infected individuals with and without HIV-1, and with and without KS. This is because in each of the four groups stratified according to these criteria, we observe all types of shedding.^34^ This observation does not preclude the possibility that oral shedding could be a valuable surrogate for the development of KS, as well as treatment response in established KS. More work is required to establish whether any metric of HHV-8 shedding can be used as a surrogate marker for risk of progression to cancer or to monitor treatment response.

The major limitation of our work is that the model is very basic. We do not ascribe a cellular source to viral reactivation from latency, for viral lytic replication, or for the acquired immune response. There is considerable knowledge about molecular mechanisms dictating the balance between HHV-8 latency and reactivation in an infected cell.^40^ Our model suggests that drugs that either sustain latency or eliminate the source of HHV-8 reactivation could be of high therapeutic value, but a major priority is to establish the nature and site of these cells in vivo. Similarly, our model attributes high importance to peripheral immunity in determining viral load but does not discriminate whether antibodies, T cells, or some combination mediate this effect. This will be important to discern to help guide the use of immunotherapy in KS and other HHV-8 related diseases.

In summary, we have created the first model of HHV-8 shedding and it performs well in terms of reproducing detailed shedding kinetics across a broad range of shedders. Future work is needed to inform mechanisms of viral latency and immune control to allow more detailed modeling.

## Acknowledgements

We are grateful to our study participants.

